# Sparse ordinal logistic regression and its application to brain decoding

**DOI:** 10.1101/238758

**Authors:** Emi Satake, Kei Majima, Shuntaro C Aoki, Yukiyasu Kamitani

## Abstract

Brain decoding with multivariate classification and regression has provided a powerful framework for characterizing information encoded in population neural activity. Classification and regression models are respectively used to predict discrete and continuous variables of interest. However, cognitive and behavioral parameters that we wish to decode are often ordinal variables whose values are discrete but ordered, such as subjective ratings. To date, there is no established method of predicting ordinal variables in brain decoding. In this study, we present a new algorithm, sparse ordinal logistic regression (SOLR), that combines ordinal logistic regression with Bayesian sparse weight estimation. We found that, in both simulation and analyses using real functional magnetic resonance imaging data, SOLR outperformed ordinal logistic regression with non-sparse regularization, indicating that sparseness leads to better decoding performance. SOLR also outperformed classification and linear regression models with the same type of sparseness, indicating the advantage of the modeling tailored to ordinal outputs. Our results suggest that SOLR provides a principled and effective method of decoding ordinal variables.

## 1 Introduction

Application of multivariate classification and regression models to functional magnetic resonance imaging (fMRI) signals has allowed the extraction of information encoded in population neural activity. Classification models are used to predict categorical variables, such as discrete stimuli and task conditions (Haynes and Rees, 2006; Norman et al., 2006; Pereira et al., 2009), while regression models are used to predict continuous parameters of interest (Cohen et al., 2011). These two types of prediction model are employed depending on the type of variable we wish to decode.

However, variables we attempt to decode are often ordinal—discrete variables whose values (classes) are ordered. For example, behavioral ratings that quantify subjective states such as the emotional feeling, impression, and preference (e.g., Baucom et al., 2012; Chang et al., 2015; Chu et al., 2011; Valente et al., 2011; Smith et al., 2014) are discrete and ordered. The intervals between classes are not defined in many cases. Furthermore, parameters of stimuli used in experiments have often been restricted to take discrete values (e.g., Kamitani and Tong, 2005, 2006; Miyawaki et al., 2008; Nishio et al., 2012; Staeren et al., 2009). Even when a parameter is defined in a metric space, the distributions of the voxel patterns are not necessarily proportionally spaced between the discrete values. Thus, discretized parameters could be better treated as ordinal variables.

Such variables have been predicted with classification and regression models in previous decoding studies. In studies using classification models, the given discrete levels were treated as nominal classes and classification models were trained to classify input brain activity patterns into one of those classes (e.g., Miyawaki et al., 2008). In studies using regression models, models were trained by treating a given ordinal variable as a continuous variable, and continuous outputs from the models were then used as the prediction results (e.g., Chang et al., 2015; Chu et al., 2011; Nishio et al., 2012; Valente et al., 2011).

Owing to the nature of ordinal variables, classification and regression models are not considered appropriate for ordinal variable prediction. An ordinal variable is a discrete variable whose classes are ordered. By definition, the distances between different classes are not given, and only the relative ordering between classes is important. In handling rating scores, for example, level 2 is placed between level 1 and level 3, but the magnitudes of the differences between levels are undefined. When a regression model is fitted using class numbers as labels, the distances between consecutive classes are treated as equal. Hence, the resultant fitness of the model depends on those deceptive distances. Meanwhile, classification models assume classes to be nominal categories and ignore given relative similarities between classes that provide helpful information for constructing a model with better prediction performance.

We here present an approach using ordinal regression, a type of generalized linear modeling whose output variable is assumed to be ordinal (Winship and Mare, 1984). In ordinal regression, similar to linear regression, a linear combination of input variables is used to predict the target variable (Figure 1A). Differently from linear regression, however, the value of the linear combination for a given input sample is not directly used as the prediction. In ordinal regression, a set of thresholds is introduced to divide the real number line into disjoint segments. These segments correspond to the discrete classes of the target variable. The class corresponding to the segment where the value of the linear combination lies is then selected as the prediction. This treats the class number as a discrete variable without using the metric in the space of the output variable. By tuning both linear weights and thresholds, ordinal regression models can be better fitted to given ordinal data than linear regression models.

**Figure 1.**
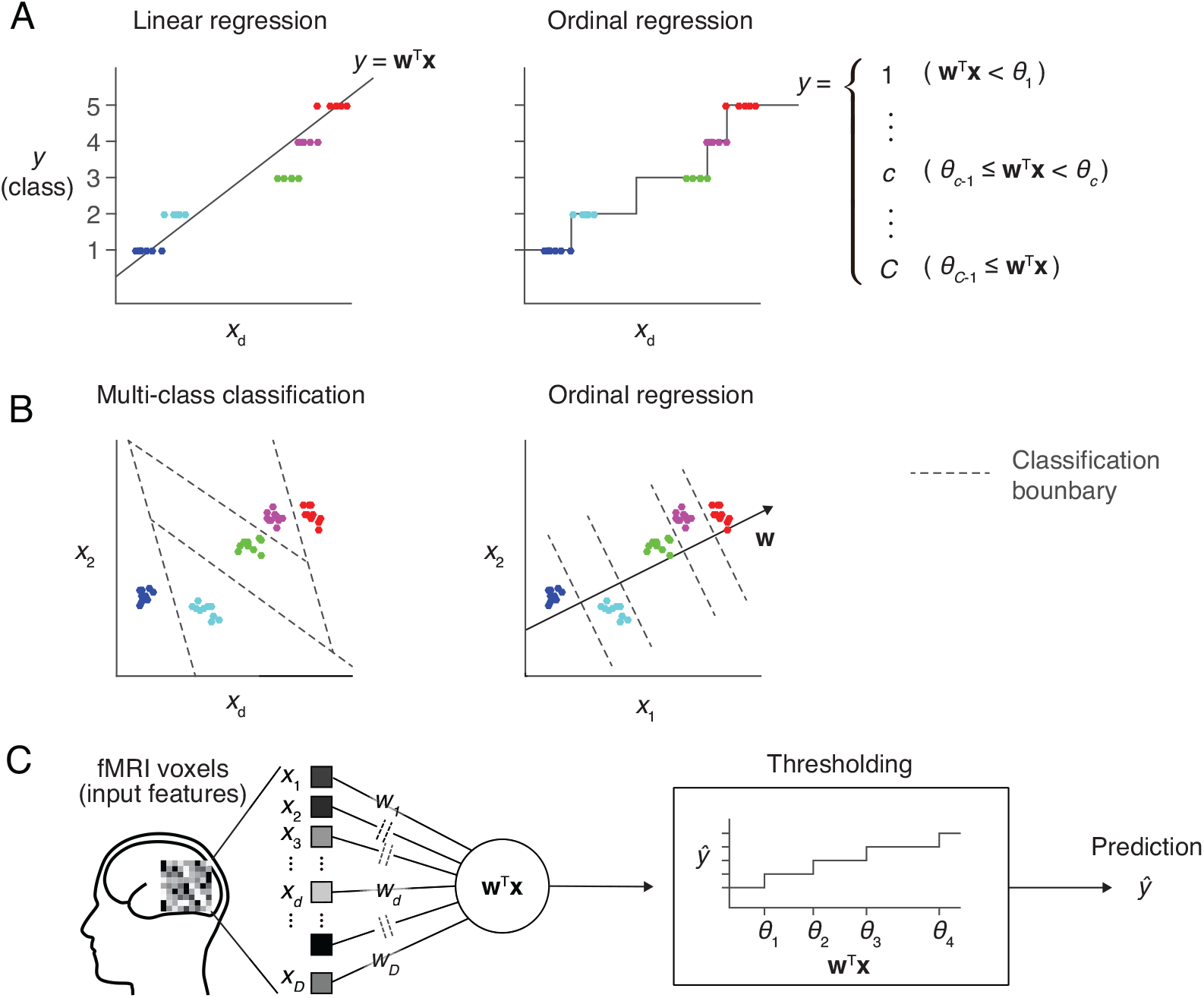
Prediction models for an ordinal dependent variable. (**A**) Comparison between linear regression and ordinal regression. In linear regression, the prediction is calculated as a linear combination of input features. Here, *y* is assumed to be an ordinal variable and it takes one of five levels. The regression line on a single input dimension (*x_d_*) is shown (left). In ordinal regression, a linear combination is used for prediction, but not directly treated as its prediction. Thresholds that divide the real number line into five disjoint segments are introduced as parameters. Each segment corresponds to one of the five levels, and the prediction is obtained as the level corresponding to the segment that the linear combination value belongs to (right). (**B**) Comparison between multiclass classification and ordinal regression. In prediction with a multiclass classification model, classification boundaries that separate different classes are learned in the feature space (left). In ordinal regression, classification boundaries are restricted to be orthogonal to one single vector, which reduces the degree of freedom compared with typical classification models (right). (**C**) Sparse ordinal logistic regression (SOLR). A standard ordinal regression model, the ordinal logistic regression model, is combined with a Bayesian sparse parameter estimation method, and is introduced into fMRI decoding analysis in the present study. By estimating a subset of weight parameters to be nonzero and the other weight parameters to be zero, this method simultaneously selects important input features (voxels) and estimates the weight parameters. In prediction, it linearly combines voxel values with the estimated sparse weight vector, then outputs one of the given levels by comparing the resultant value with thresholds.

Compared with classification models, ordinal regression models are expected to be efficient in learning, leading to better prediction performance. Classification models learn decision boundaries that are used to classify input samples into classes in the feature space (Figure 1B, left). Their degree of freedom increases as the number of classes increases, which makes parameter estimation sensitive to noise. In contrast, all decision boundaries of an ordinal regression model are restricted to be orthogonal to a single line in the feature space, and the degree of freedom is smaller than that for classification models (Figure 1B, right). This lower complexity of ordinal regression models reduces the chance of overfitting and lead to better generalization performance than classification models.

To introduce a multivariate prediction model into fMRI decoding analysis, it is generally important to choose an appropriate set of input voxels because the presence of many irrelevant voxels can lead to poor generalization performance due to overfitting. In standard decoding analysis, only tens or hundreds of fMRI samples are obtained to train the prediction model, while the input feature vector consists of thousands of voxels. Thus, overfitting readily occurs if all available voxels are used as input features. To solve this problem, our previous study proposed a classification algorithm that simultaneously performs voxel selection and parameter estimation, and demonstrated that the method successfully prevents overfitting in the presence of many irrelevant voxels (Yamashita et al., 2008). In that study, a Bayesian extension of logistic regression was proposed where the automatic relevance determination (ARD; MacKay 1992; Neal 1996) prior was used as the prior distribution of the weight vector. This resulted in selecting a small number of voxels as important by estimating the corresponding weight parameters to be nonzero, and ignoring the other voxels by estimating their weight parameters to be zero. This sparse parameter estimation provided a method of voxel selection by virtually eliminating voxels associated with zero-valued weight parameters.

In the present study, we combine ordinal regression with the sparse estimation (Figure 1C) to build an ordinal prediction model suited to fMRI decoding. As our model is based on ordinal logistic regression (OLR; McCullagh, 1980), a standard ordinal regression model, we refer to our proposed method as sparse ordinal logistic regression (SOLR). We evaluate the performance of SOLR using both simulation and real fMRI data. In these analyses, the prediction performance of SOLR is compared with that of an OLR model without a sparseness constraint to examine the utility of the sparseness. Likewise, the prediction performance is compared with that of regression and classification models having the same type of sparseness, sparse linear regression (SLiR; Bishop, 2006; Tipping, 2001) and sparse multinomial logistic regression (SMLR; Yamashita et al., 2008), to examine the superiority of SOLR in ordinal variable prediction. To examine whether SOLR works well in practical situations, we compare the decoding performances for different numbers of training samples and input dimensions in the simulation analysis. In the analysis using real fMRI data, we tested the four previously mentioned algorithms on a dataset taken from Miyawaki et al. (2008). In this previous study, stimulus images were reconstructed from fMRI responses by training decoders on 440 samples using about 1000 voxels from the primary visual area (V1) as input. Using this dataset, we demonstrate that SOLR better predicts ordinal variables in a practical situation of fMRI decoding analysis.

## 2 Materials and Methods

### 2.1 Algorithm

This section first describes OLR, which is a generalized linear model for ordinal dependent variables (Winship and Mare, 1984; McCullagh, 1980), and then explains SOLR by introducing a Bayesian framework to estimate parameters. OLR with L2-regularization (L2OLR) is also explained in this section. Our MATLAB and Python implementations of SOLR and L2OLR are available at our Github respository^1^.

OLR is one of the generalized linear models whose dependent variable is assumed to be an ordinal variable. In OLR, the dependent variable 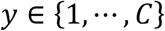 is assumed to follow the underlying process given by

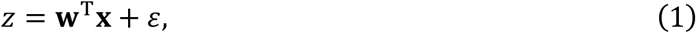

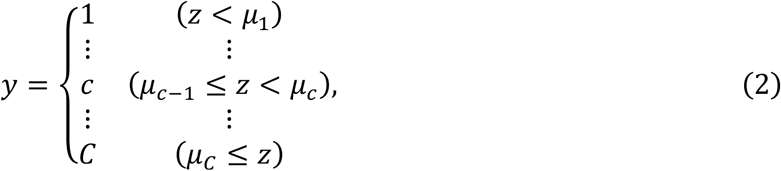

where 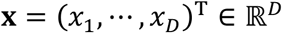 is the vector of *D* independent variables (input features), 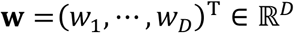 is the linear weight vector, *ε* is a random variable representing the noise, *z* is a latent variable in the model, and *μ*_1_, *μ*_2_, ⋯, *μ*_*C*−1_ (*μ*_1_ ≤ *μ*_2_ ≤ ⋯ ≤ *μ*_*C*−1_) are threshold parameters. In OLR, *ε* is assumed to follow the logistic distribution with a mean of zero and a variance of 1. The threshold parameters are collectively denoted by a single vector **μ**.

In OLR, **w** and **μ** are estimated by maximizing the log likelihood function with a gradient method. The log likelihood with respect to **w** and **μ** is given by

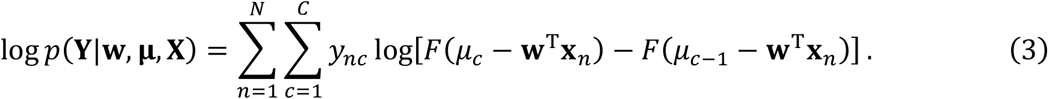

Here, *y_nc_* is a binary variable indicating whether the value of the dependent variable for the *n*-th sample is *c. y_nc_* is set to 1 if the value of the dependent variable for the *n*-th sample is *c* and *y_nc_* is set to zero otherwise (1-of-k representation). **x**_*n*_ is the vector of feature values for the *n*-th sample. *N* samples are used for estimation, and are collectively denoted by the *N* × *C* matrix **Y** and the *N* × *D* matrix **X**. The function *F* is the logistic sigmoid function defined by

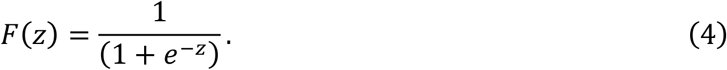

In the above log likelihood function, the first and last threshold parameters *μ*_0_ and *μ_C_* are respectively set to −∞ and +∞ by convention.

We next introduce a Bayesian framework to estimate the above parameters sparsely. We introduce prior distributions for parameters to be estimated in OLR. For the parameter **w**, we assume that

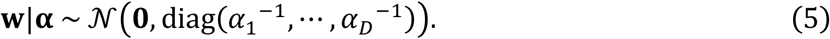

Here *α*_1_,⋯, *α_D_* are hyperparameters that determine the importance of voxels and are called relevance parameters. They are collectively denoted by the single vector 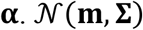 represents the multidimensional Gaussian distribution with mean **m** and covariance **Σ. 0** represents the zero vector, while diag(*α*_1_^−1^,⋯, *α*_*D*_^−1^) represents the diagonal matrix whose diagonal elements are *α*_1_^−1^,⋯, *α*_*D*_^−1^ and nondiagonal elements are zero. If *α*_*d*_^−1^ is small, the distribution function of *W_d_* has a sharp peak around zero and the corresponding voxel thus tends to be virtually ignored in prediction. Meanwhile, if *α*_*d*_^−1^ is large

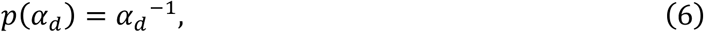

as often adopted in previous studies (Yamashita et al., 2008). Additionally, for the parameter **μ**, we assume the noninformative prior, which is expressed as

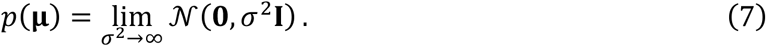

We then obtain the log of the posterior distribution as

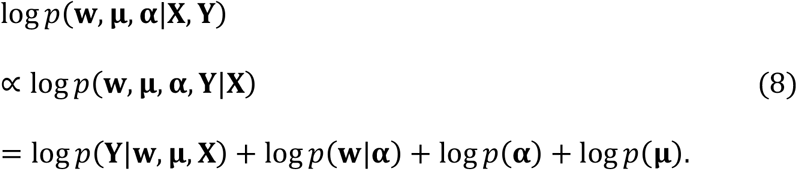

In the present study, the values of **w, μ**, and **α** that maximize the above function—the maximum a posteriori (MAP) solution—were estimated with training data, and the values of **w** and **μ** were then used in prediction on test data. Because the MAP solution for the above cannot be derived in a closed form, we used the mean-field variational Bayesian approximation and the Laplace approximation (Attias 1999, Bishop, 2006; see Appendix). Once we obtain the MAP solution, we can calculate the predictive probability of each class for a given new input vector. The class with the highest predictive probability was chosen as the prediction outcome.

To examine the effect of voxel selection by ARD, we compared the performance of SOLR with that of OLR having L2-regularization (L2OLR). In L2OLR, we assume that the prior distribution of **w** is the Gaussian distribution with zero mean and isotropic covariance, as expressed by

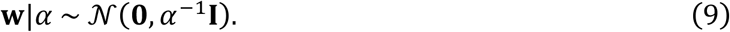

*α* is a hyperparameter that controls the degree of regularization. In a similar manner to SOLR, we assume noninformative priors for *α* and **μ**. We estimated the MAP solution with the same approximation techniques, and then made a prediction using the estimated parameters.

### 2.2 Simulation analysis

We compared the prediction performance across SOLR, L2OLR, SLiR, and SMLR using simulation data. For data generation, five *D*-dimensional Gaussian distributions were prepared, and samples for class *c* were generated from the *c*-th Gaussian distribution. The means of the Gaussian distributions were given by

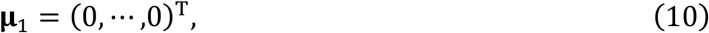

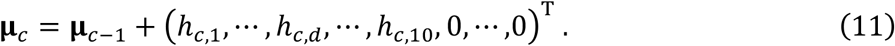

*h_c,d_* (*c* = 2, ⋯ 5; *d* = 1, ⋯, 10) are parameters that specify the intervals between the means, and each was sampled from an exponential distribution with a mean of 1.0. Only the first 10 dimensions have information on classes, and the other dimensions are irrelevant. In each of the first 10 dimensions, the mean of the input feature monotonically increases against the class label, which leads to an ordinal structure in the feature space. We also conducted the same simulation analysis in the case that the means of the Gaussian distributions are equally spaced by setting all *h_cd_* to 1.0, and observed qualitatively similar comparison results. The covariance matrices of the Gaussian distributions were set as diagonal matrices regardless of the class label. The standard deviation in each dimension was set to 3.0. *D* was set to 10, 50, 100, 250, 500, 1000, 1500, and 2000 to characterize the prediction performance as a function of the number of input dimensions.

To evaluate the prediction performance, a prediction model was trained and tested on independent sets of samples. *N* samples and 1000 samples were respectively generated from the same Gaussian distributions as training and test data. *N* was set to 10, 25, 50, 100, 250, 500, 1000, 1500, and 2000 to characterize the prediction performance as a function of the number of training samples. Equal numbers of samples were generated for the five classes. To quantify the prediction performance, we calculated the Spearman rank correlation between true and predicted labels using test data. The same simulation procedure was repeated 100 times, and the prediction performance measured by the Spearman rank correlation was averaged across those 100 repetitions.

### 2.3 fMRI data analysis

We compared the prediction performance across the four algorithms using real fMRI data from Miyawaki et al. (2008). The dataset can be downloaded from public databases^2^ (Poldrack et al., 2013; Takemiya et al., 2016). It contains fMRI signals when the subject was viewing visual images consisting of contrast-defined 10 × 10 checkerboard patches. Each patch was either a flickering checkerboard or a homogeneous gray area. The dataset consists of two independent sessions. One is a random image session, in which a spatially random pattern was presented for 6 s and there was a subsequent 6-s rest period. A total of 440 different random patterns were presented to the subject. The other is a figure image session, where a letter of the alphabet or a simple geometric shape was presented for 12 s and there was a subsequent 12-s rest period. Five letters of the alphabet and five geometric shapes were presented eight times.

Miyawaki et al. (2008) successfully reconstructed presented images from fMRI responses by combining multiple classifiers. To reconstruct arbitrary visual images, a set of local regions that cover the entire stimulus image area was predefined, the mean contrast in each local region was then predicted by a classifier, and the outputs from the classifiers for those local regions were then optimally combined to produce a single reconstructed image. In the previous study, SMLR was used to construct classifiers. Here, we used SOLR, L2OLR, SLiR, and SMLR for contrast prediction, and compared the prediction performance among them.

## 3 Results

### 3.1 simulation analysis

In the simulation analysis, samples from the five classes were generated from five multidimensional Gaussian distributions, and a prediction model using each algorithm was trained and tested on independent sets of data samples (see Materials and Methods). We set only the first 10 dimensions to have information on the classes while keeping the other dimensions irrelevant. In each of the first 10 dimensions, the centers of the five Gaussian distributions were placed so that the mean of the input feature monotonically increases against the class number to assume an ordinal structure in the feature space.

To characterize prediction performance when the number of input dimensions is large, performance was calculated as a function of the number of input dimensions (Figure 2A). Prediction performance was evaluated using the Spearman rank correlation between true and predicted labels. The number of training samples was fixed to 100, which is a typical size in real fMRI decoding analysis. While all algorithms had similar performance when the number of input dimensions was small, SOLR outperformed the other algorithms as the number increased.

**Figure 2.**
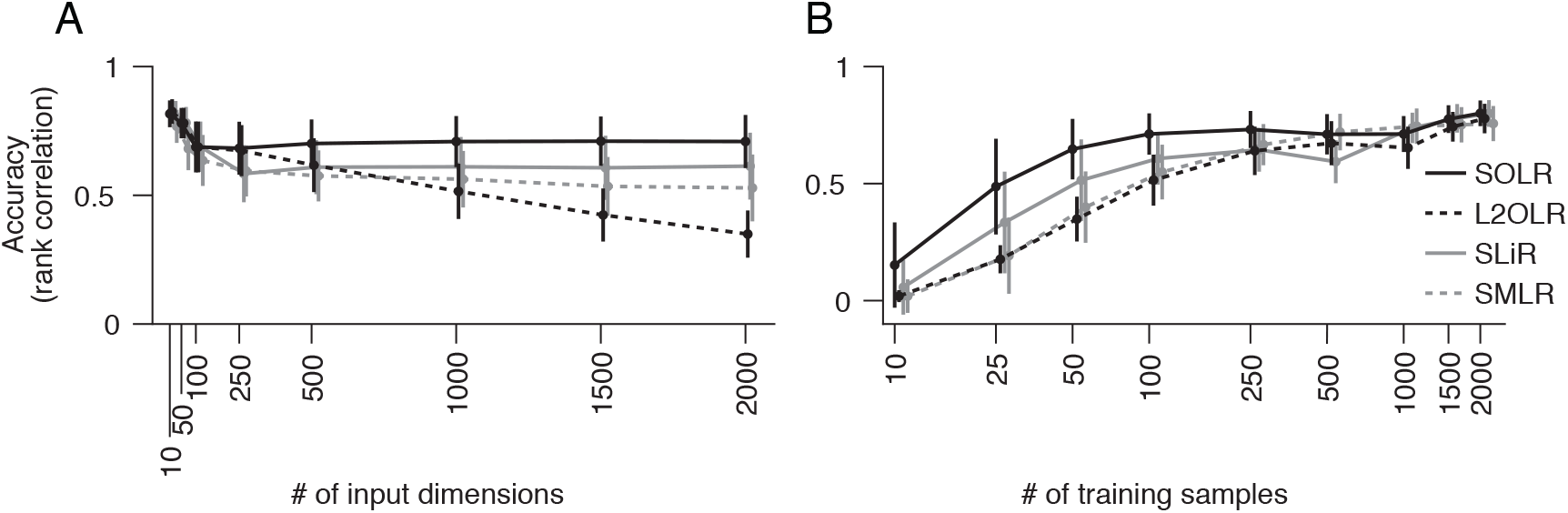
Simulation analysis. (**A**) Decoding accuracy as a function of the number of input dimensions. SOLR, L2OLR, SLiR, and SMLR are trained with 100 samples and tested with 1000 samples using simulation data. The prediction performance is evaluated by the Spearman rank correlation between true and predicted labels. In this simulation, only 10 input dimensions include information on labels, and the number of irrelevant dimensions is changed to characterize the performance. The mean performance across 100 simulation repetitions is plotted as a function of the total number of input dimensions. Error bars show the standard deviations across 100 simulation repetitions. (**B**) Decoding accuracy as a function of the number of training data. The mean prediction performance is plotted as a function of the number of training samples. The total number of dimensions is fixed to 1000. Error bars show the standard deviations across 100 simulation repetitions.

Furthermore, the prediction performance was calculated as a function of the number of training samples to characterize the performance when the number of training data is small (Figure 2B). Here, the number of input dimensions was fixed to 1000, which is a typical input size in decoding analysis. As a result, SOLR had higher performance than the other algorithms when the number of training data was small. As we increased the number of training samples, all algorithms reached similar accuracies.

### 3.2 fMRI data analysis

We also evaluated the prediction performance using the real fMRI dataset from Miyawaki et al. (2008). The cited study measured fMRI responses as the subject viewed 10 × 10 binary images and successfully reconstructed arbitrary visual images from the fMRI responses (Figure 3A,B). In the reconstruction procedure, it was assumed that an image can be represented by a linear combination of local image bases of multiple scales (1 × 1, 1 × 2, 2 × 1, and 2 × 2; Figure 3A). There are a few possible stimulus states (contrast patterns) in the region specified by a single image basis, and the stimulus states can be classified according to the mean contrasts (Figure 3C). The mean contrast for the 1 × 1 image basis is binary, but the mean contrast for 1 × 2 and 2 × 1 image bases takes one out of three discrete values, and that for the 2 × 2 image basis takes one out of five discrete values. While the intervals between the successive mean contrasts can be regarded as equal in the contrast space, the distributions of voxel patterns for the discrete values are not necessarily equally spaced. In fact, the amplitude of the most responsive voxel for each image basis monotonically increases against but is not proportional to the mean contrast level (Figure 3D). Miyawaki et al. (2008) predicted the mean contrasts for the image bases using binary or multiclass classifiers based on sparse logistic regression (sparse [multinomial] logistic regression, SMLR), disregarding the order of contrasts. Here, we used SOLR, L2OLR, SLiR, and SMLR to predict the mean contrasts for 1 × 2, 2 × 1, and 2 × 2 image bases, and compared the prediction performance among them.

**Figure 3.**
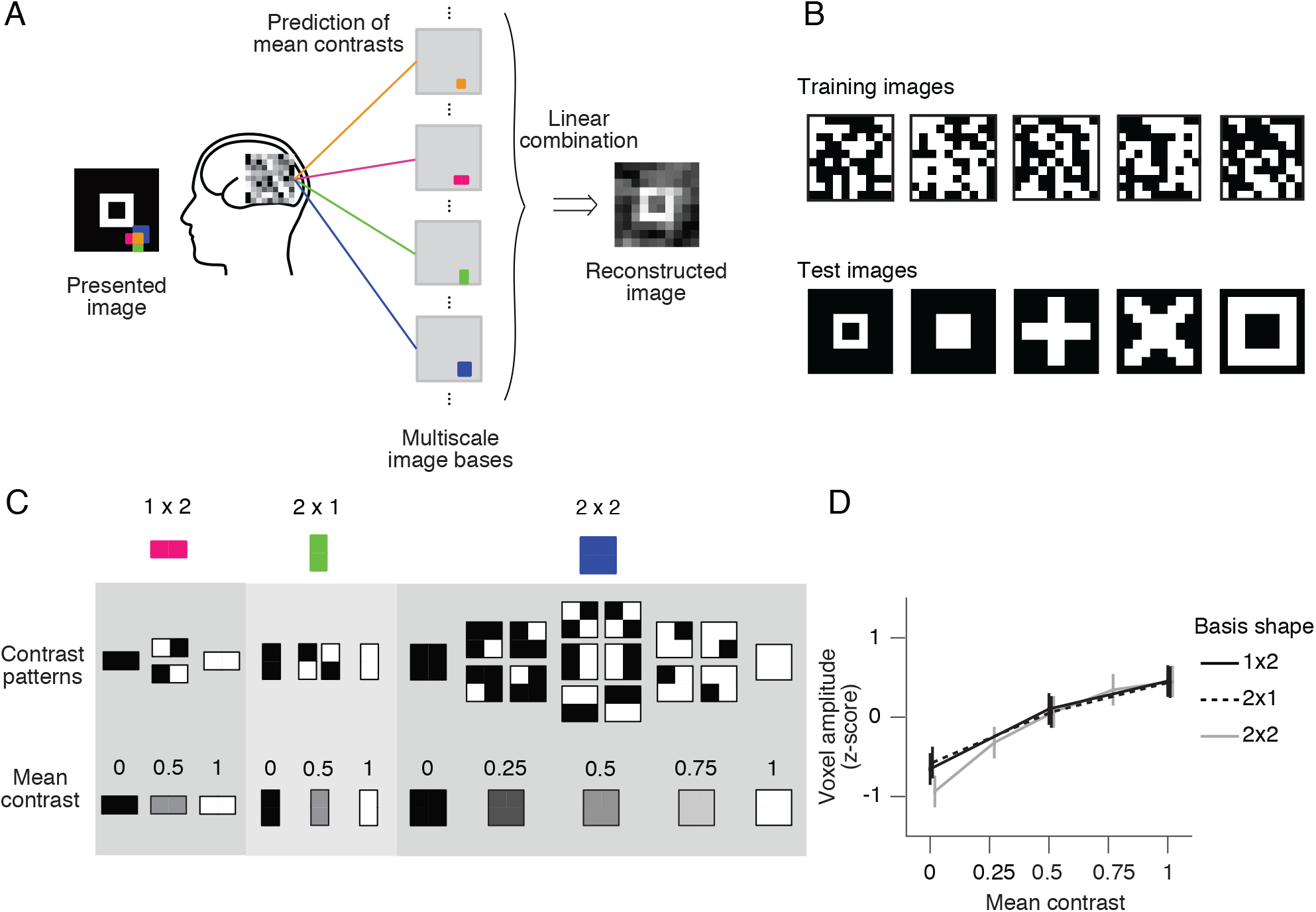
Ordinal regression task with visual image reconstruction data. (**A**) Reconstruction procedure of Miyawaki et al (2008). A presented image was reconstructed by first predicting the contrasts of image bases and then optimally combining the image bases multiplied by the contrasts. Image bases were 1 × 1-, 1 × 2-, 2 × 1-, and 2 × 2-shaped regions covering a 10 × 10-pixel image with overlaps. (**B**) Examples of stimulus images. Random binary images (upper) and simple geometric shapes (lower) were respectively presented in the random image session and the figure image session of the experiment. In reconstruction analysis, data in the random image session were used to train decoders and data in the figure image session were used as test data. (**C**) Contrast patterns and mean contrast in image bases. 1 × 2- and 2 × 1-shaped regions took one of the four contrast patterns, and the mean contrast within the region took one of the three values (left, middle). 2 × 2-shaped regions took one of the 16 contrast patterns, and the mean contrast took one of the five values. (**D**) Relationship between the voxel amplitude and mean contrast. Voxels responsive to each local region were selected in a univariate analysis of the random image session data. The voxel amplitudes were then averaged across trials for each mean contrast. Error bars show the 95% confidence intervals across local regions.

We evaluated the prediction performance of each algorithm by five-fold cross-validation using the random session data (Figure 4A). The prediction performance for each image basis was quantified by the Spearman rank correlation between true and predicted contrasts across test samples. The continuous outputs of SLiR were assigned to the nearest discrete labels. For all basis shapes, SOLR outperformed the other algorithms. The median performances of SOLR were significantly higher than those of L2OLR (*p* < 0.001, signed-rank test), SLiR (*p* < 0.001), and SMLR (*p* < 0.005) for all basis shapes.

**Figure 4.**
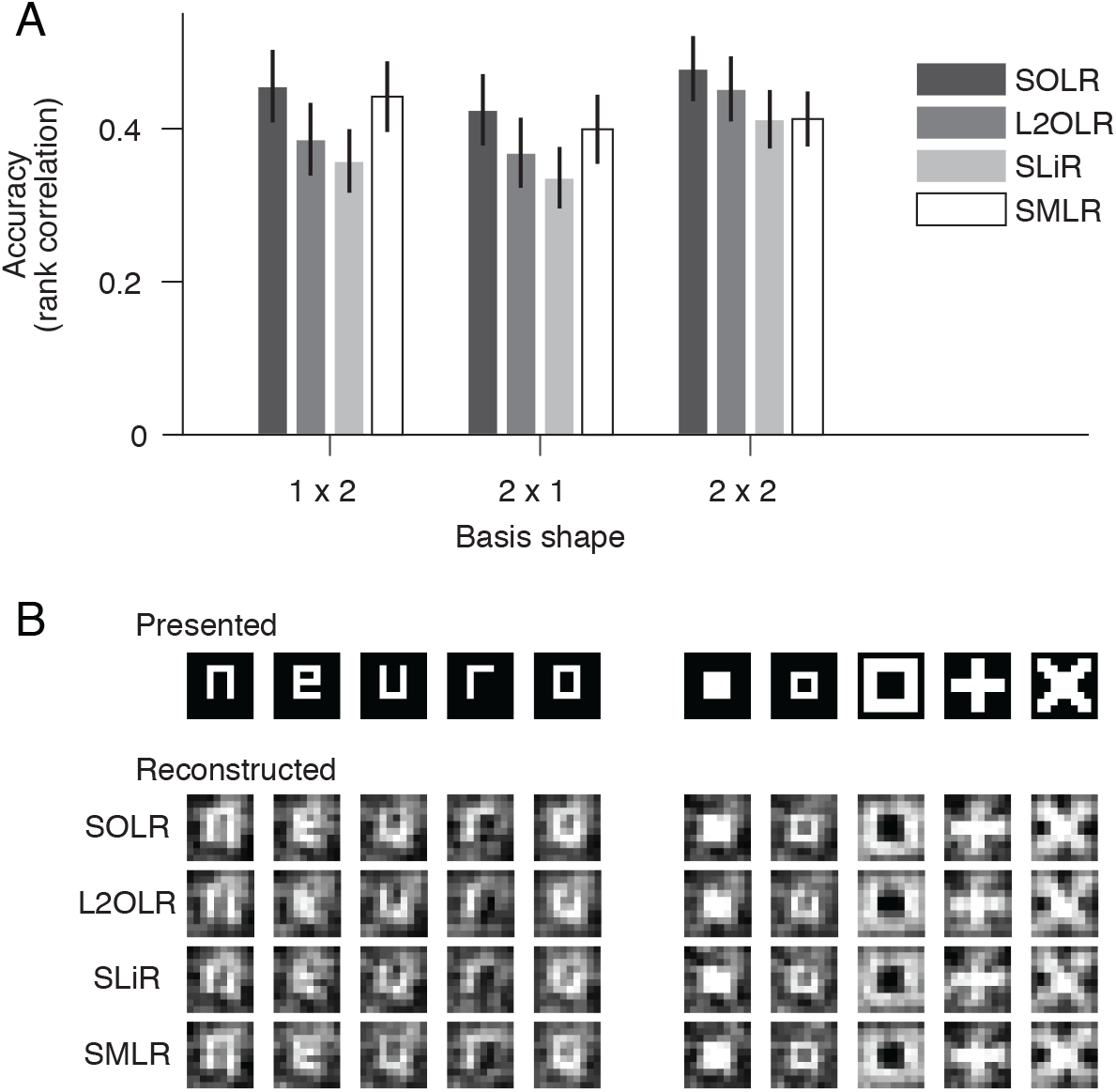
Performance on visual image reconstruction data. (**A**) Prediction of mean contrasts of image bases. For each basis, the mean contrast was predicted by SOLR, L2OLR, SLiR, and SMLR. Performance was evaluated using the Spearman rank correlation between true and predicted mean contrast values across test samples. The mean performance averaged across basis locations is shown. Error bars show the 95% confidence intervals across basis locations. (**B**) Reconstructed images. Presented images and images reconstructed with the four algorithms are shown.

We finally reconstructed visual images according to the predictions of the models with the same procedure as adopted by Miyawaki et al. (2008), and compared reconstructed images between the four algorithms (Figure 4B). The image bases were multiplied by the predicted mean contrasts and then linearly combined with optimized weights to produce a single image (see Miyawaki et al., 2008 for details). Although the differences are not remarkable in visual inspection, the spatial correlations between presented images and images reconstructed with SOLR were higher than those of the other algorithms (*p* < 0.05, signed-rank test).

## 4 Discussion

We developed a new algorithm for ordinal variable decoding by combining OLR with Bayesian sparse weight estimation. The proposed algorithm, SOLR, was compared with three other methods: (1) ordinal logistic regression without a sparse constraint, L2OLR; (2) a regression model with the same Bayesian sparse constraint, SLiR; and (3) a classification model with the same Bayesian sparse constraint, SMLR. In analyses using simulation and real fMRI data, SOLR had better prediction performance than the other three methods. These results suggest that SOLR is a useful tool in decoding analyses where the target variable can be regarded as ordinal.

Ordinal variables naturally emerge in decoding analysis; however, they have been predicted using classification models (Baucom et al., 2012; Cortese et al., 2016, 2017; Miyawaki et al., 2008; Staeren et al., 2009) or regression models (Chang et al., 2015; Chu et al., 2011; Nishio et al., 2012; Valente et al., 2011). By the nature of the ordinal variable, the levels an ordinal variable takes have a relative order but the distances between levels are not given. Because regression models use a metric in the label space and their predictions depend on it (Figure 1A), regression models are not appropriate for ordinal variable prediction. Meanwhile, classification models do not need the distances between classes. However, the complexity of classification models rapidly grows as the number of classes becomes large, which increases the chance of overfitting (Figure 1B). Here, to predict ordinal variables in decoding analysis, we introduced OLR (McCullagh, 1980), one of the known generalized linear models whose output variable is assumed to be an ordinal variable. To prevent overfitting in decoding analysis where a large number of voxels are used as input, we proposed a new method, SOLR, by combining OLR with a Bayesian sparse weight estimation method (MacKay 1992; Neal 1996; Yamashita et al., 2008).

In the analysis using simulation data, SOLR outperformed L2OLR, SLiR, and SMLR as the number of input dimensions increased or the number of training data decreased (Figure 2). The comparison between SOLR and L2OLR suggests that the sparseness introduced into SOLR prevents overfitting efficiently and improves the decoding performance, which is consistent with the results of previous studies analyzing the utility of the sparseness using classification models (Ryali et al., 2010; Yamashita et al., 2008). A comparison among SOLR, SLiR, and SMLR showed that the appropriate treatment of a given relative order by OLR also leads to better decoding performance.

In analysis using real fMRI data, the same four methods were compared and SOLR had better prediction performance than L2OLR, SLiR, and SMLR (Figure 4A). While the same contrast prediction task was conducted with SMLR in the previous study (Miyawaki et al., 2008), we found that the prediction can be improved by introducing SOLR. Although the resultant reconstructed images of SOLR and SMLR appear similar (Figure 4B), SOLR had a slightly higher spatial correlation than SMLR. These results suggest that SOLR would work well in a practical situation of fMRI decoding analysis.

Taken together, SOLR is expected to provide a principled and effective method of decoding ordinal variables. While ordinal variables have been predicted using classification models or regression models in previous decoding studies, we found that SOLR outperformed linear classification and regression models with the same type of sparseness. The results suggest that SOLR would be helpful in decoding analysis where an ordinal variable is used as the target variable and would allow us to better characterize the neural representations of subjective states that are quantified by subjective ratings, such as impressions, emotional feelings, and confidence.

## Author Contributions

ES, KM, and YK designed the study. KM developed the algorithm. ES and SCA performed analyses. ES, KM, and YK wrote the manuscript.

## Acknowledgments

The authors thank Mohamed Abdelhack, Tomoyasu Horikawa, Ken Shirakawa, Yu Takagi, Mitsuaki Tsukamoto, and the members of Kamitani Lab at Kyoto University and ATR computational neuroscience laboratories for helpful comments on the manuscript and analysis.

## Appendix

We estimated the MAP solution of the weight and threshold parameters in SOLR using the mean-field variational Bayesian approximation and Laplace approximation. In this estimation procedure, we approximate the posterior distribution as

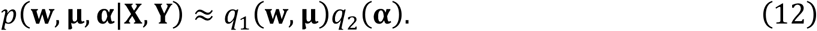

*q*_1_ and *q*_2_ are probability density functions that are iteratively updated to obtain a better approximation. In the variational Bayesian method, *q*_1_ and *q*_2_ are alternately updated using the rules

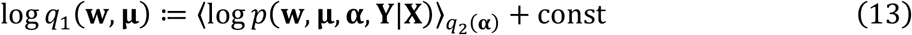

and

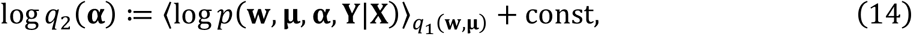

where 〈*x*〉_*q*(*x*)_ denotes the expectation of *x* with respect to the probability distribution *q*(*x*). Each update decreases the Kullback–Leibler divergence between *p*(**w, μ, α|X, Y**) and *q*_1_(**w, μ**)*q*_2_(**α**), which makes *q*_1_(**w, μ**)*q*_2_(**α**) a better approximation of the posterior distribution (Attias, 1999; Yamashita et al., 2008; Bishop, 2006). In the following, we describe the procedure of updating *q*_1_ and *q*_2_, respectively.

To update *q*_1_, the right side of (13) is rewritten as

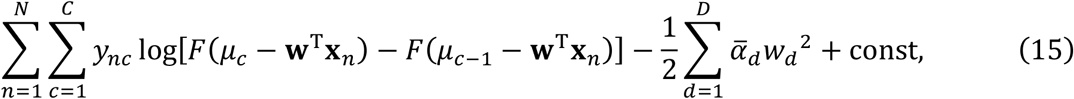

where 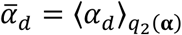. *F* is the logistic sigmoid function whose definition was given in (4). The probability distribution function whose logarithm is given by (15) cannot be obtained in analytic form, and we therefore applied the Laplace approximation. In the approximation, (15) is replaced with its second-order Taylor series expansion around the maximum, and the update equation of *q*_1_ is rewritten as

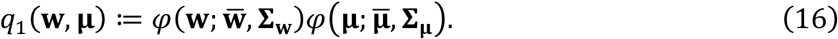

Here, the function *φ*(·; **m, Σ**) denotes the probability density function of the multidimensional Gaussian distribution with mean **m** and covariance **Σ**. 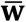 and 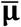 are the values of **w** and **μ** that maximize 〈log*p*(**w, μ, α, Y|X**)〉_*q*_2_(**α**)_. **Σ_w_** is the Hessian matrix of 〈log*p*(**w, μ, α, Y|X**)〉_*q*_2_(**α**)_ with respect to **w** at 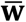. **Σ_μ_** is the Hessian matrix of 〈log *p*(**w, μ, α, Y|X**)〉_*q*_2_(**α**)_ with respect to **μ** at 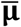. The calculation of 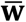 and 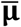 was performed using a gradient method in the present study. As **Σ_μ_** is not used in the update of *q*_2_, only 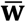, 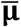, and **Σ_w_** were calculated in the present study.

In the update of *q*_2_, using the relationship (16), the update rule is given by

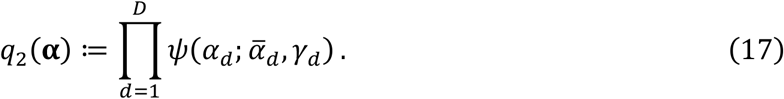

Here, 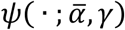 denotes the probability density function of the gamma distribution with mean 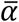 and degree of freedom *γ. γ_d_* is 0.5 regardless of *d*, and 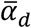 is given by

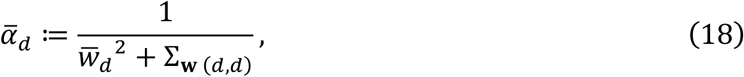

where 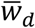 is the *d*-th element of **w** and Σ_**w**(*d,d*)_ is the *d*-th diagonal element of **Σ_w_**. To accelerate convergence, a modified rule adopted in previous studies (MacKay, 1992; Yamashita et al., 2008) was used instead of (18):

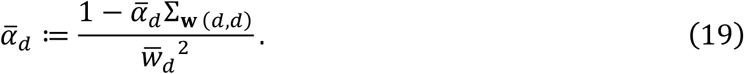

As initial parameters, 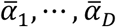, were set to 1, *q*_1_ and and *q*_2_ were alternately updated 100 times in this study.

## Footnotes

1 https://github.com/KamitaniLab/SOLR

2 https://openfmri.org/ http://brainliner.jp/data/brainliner/Visual_Image_Reconstruction

